# qPRF: A system to accelerate population receptive field decoding

**DOI:** 10.1101/2024.08.13.607805

**Authors:** Sebastian Waz, Yalin Wang, Zhong-Lin Lu

## Abstract

Patterns of BOLD response can be decoded using the population receptive field (PRF) model to reveal how visual input is represented on the cortex (Dumoulin and Wandell, 2008). The time cost of evaluating the PRF model is high, often requiring days to decode BOLD signals for a small cohort of subjects. We introduce the qPRF, an efficient method for decoding that reduced the computation time by a factor of 1436 when compared to another widely available PRF decoder (Kay, Winawer, Mezer and Wandell, 2013) on a benchmark of data from the Human Connectome Project (HCP; Van Essen, Smith, Barch, Behrens, Yacoub and Ugurbil, 2013). With a specially designed data structure and an efficient search algorithm, the qPRF optimizes the five PRF model parameters according to a least-squares criterion. To verify the accuracy of the qPRF solutions, we compared them to those provided by Benson, Jamison, Arcaro, Vu, Glasser, Coalson, Van Essen, Yacoub, Ugurbil, Winawer and Kay (2018). Both hemispheres of the 181 subjects in the HCP data set (a total of 10,753,572 vertices, each with a unique BOLD time series of 1800 frames) were decoded by qPRF in 15.2 hours on an ordinary CPU. The absolute difference in *R*^2^ reported by Benson et al. and achieved by the qPRF was negligible, with a median of 0.39% (*R*^2^ units being between 0% and 100%). In general, the qPRF yielded a slightly better fitting solution, achieving a greater *R*^2^ on 99.7% of vertices. The qPRF may facilitate the development and computation of more elaborate models based on the PRF framework, as well as the exploration of novel clinical applications.

**Highlights:**

- We describe a novel software system, qPRF, which can perform population receptive field (PRF) decoding of BOLD fMRI at speeds about 1400 times faster than the conventional systems designed for PRF decoding.
- We show that qPRF yields estimates of PRF model parameters that, in terms of goodness-of-fit, are equivalent to estimates derived using the conventional systems.
- An efficient similarity-based search strategy, underlies the accelerated computations of qPRF, supported by a specially designed data structure wherein tens of millions of pre-computed prediction curves are stored.

## 1. Introduction

Retinotopic maps – representations of the visual field on the surface of the brain – have been a subject of scientific inquiry for at least a century (Ribeiro, Benson and Puckett, 2024), established in large part initially by studies of brain lesions suffered by people injured in war (Glickstein and Whitteridge, 1987; Inouye, 1909; Holmes and Lister, 1916) and subsequently by surgical investigations in homologues. Physiological measurements of retinotopic maps in humans were made as early as the 1980s using positron emission topography (Fox, Mintun, Raichle, Miezin, Allman and Van Essen, 1986; Fox, Miezin, Allman, Van Essen and Raichle, 1987), and in the following decade, the current conventions for studying human retinotopic maps began to emerge (DeYoe, Carman, Bandettini, Glickman, Wieser, Cox, Miller and Neitz, 1996; Dumoulin, Hoge, Baker Jr, Hess, Achtman and Evans, 2003; Engel, Rumelhart, Wandell, Lee, Glover, Chichilnisky, Shadlen et al., 1994; Engel, Glover and Wandell, 1997; Sereno, Dale, Reppas, Kwong, Belliveau, Brady, Rosen and Tootell, 1995), namely, the use of functional magnetic resonance imaging (fMRI) to detect blood-oxygenation-level-dependent (BOLD) signals that were phase-locked to visual stimulation. Based on these developments, Dumoulin and Wandell (2008) formulated the population receptive field (PRF) model which has become the standard model in retinotopic mapping for decoding BOLD signals. Here, we present a novel algorithm, qPRF, to accelerate population receptive field decoding.

The PRF model has had a singular influence within visual neuroscience. For example, the current largest freely available retinotopy dataset, collected by the Human Connectome Project (HCP; Uğurbil, Xu, Auerbach, Moeller, Vu, Duarte-Carvajalino, Lenglet, Wu, Schmitter, Van de Moortele, Strupp, Sapiro, De Martino, Wang, Harel, Garwood, Chen, Feinberg, Smith, Miller, Sotiropoulos, Jbabdi, Andersson, Behrens, Glasser, Van Essen and Yacoub, 2013; Van Essen et al., 2013), is based on an experiment designed specifically to maximize the test-retest reliability of PRF estimates; in fact, for many researchers within this area, their main interface with the HCP retinotopy dataset is not the BOLD time series which comprise the raw data, but the corresponding PRF model estimates provided by Benson et al. (2018) for this dataset. Currently, the PRF model represents the gold standard for processing BOLD fMRI in the course of identifying the layout and boundaries of visual areas (e.g., Dougherty, Koch, Brewer, Fischer, Modersitzki and Wandell, 2003, Benson, Yoon, Forenzo, Engel, Kay and Winawer, 2022, Himmelberg, Tünçok, Gomez, Grill-Spector, Carrasco and Winawer, 2023) which, subsequently, enables neuroscientists to study the functional specialization of the regions of the brain (Silver and Kastner, 2009). Given its unique importance within visual neuroscience, the original article of Dumoulin and Wandell (2008) has received more than 1,000 citations according to Google Scholar, and according to Scopus, the paper is in the 96^th^ percentile compared to similar papers in terms of citation frequency. Elaborations on the PRF model are being actively developed, with a number of iterations already described in the literature (e.g., Kay et al., 2013; Haak, Winawer, Harvey, Renken, Dumoulin, Wandell and Cornelissen, 2013; Zeidman, Silson, Schwarzkopf, Baker and Penny, 2018).

In addition to its scientific influence, the output from PRF modeling shows great promise as an endpoint in clinical applications. In Alzheimer’s disease (AD), visual deficits are commonly reported as an early symptom, and these deficits have been associated with reduced surface area of the retinotopic maps in the brains of patients with AD relative to controls (Brewer and Barton, 2014; Brewer, Barton, Brewer and Barton, 2016). Glaucoma, a disease which leads to damage of the optic nerve, can be assessed with fMRI (Duncan, Sample, Weinreb, Bowd and Zangwill, 2007; Boucard, Hernowo, Maguire, Jansonius, Roerdink, Hooymans and Cornelissen, 2009), so PRF modeling may prove useful in tracking its progression (thus making it useful as an endpoint for medications in treating glaucoma). Although we know of no reported usage of the PRF model in neurosurgery, Hense, Plank, Wendl, Dodoo-Schittko, Bumes, Greenlee, Schmidt, Proescholdt and Rosengarth (2021) have shown that retinotopic organization can be reliably assessed in pre-operative patients with tumors and/or lesions of the occipital lobe. Similar technologies have been used in the pre-surgical planning and post-surgical assessment of patients receiving an implant for deep brain stimulation (Lauro, Lee, Ahn, Barborica and Asaad, 2018). The success of these interventions suggests that the PRF model might be useful in neurosurgery for patients dealing with visual impairments like those described above (Bridge, 2011). For those with stroke-induced cortical lesions, retinotopic maps may also be reliably estimated (Elshout, Bergsma, van den Berg and Haak, 2021), and their corresponding visual deficits may be recovered using noninvasive interventions such as visual restitution training (VRT); the improvement observed under VRT is associated with increased PRF size within the cortical representation of the blind field (Barbot, Das, Melnick, Cavanaugh, Merriam, Heeger and Huxlin, 2021). In patients suffering macular degeneration (MD), a disease marked by central retina damage (Bridge, 2011), peripheral stimulation has been found to activate the foveal confluence (Baker, Peli, Knouf and Kanwisher, 2005), suggesting passive adaptation in the visual system of those with MD that resembles the adaptation achieved by VRT.

Currently, computing PRF model fits to retinotopy fMRI data is time intensive, and this can pose a challenge to users of the model. Although the exact time requirements will depend on whether the user has access to specialized resources such as a high-performance computing cluster, for a single brain with 91,282 vertices, we estimate that it can take 194 hours to generate a complete PRF solution using a popular implementation of the model, analyzePRF (Kay et al., 2013) on a typical desktop computer with mutli-core processor (in our case, a 3.50GHz Intel Xeon E5-1650 v3 CPU). Such a time demand can restrict how the model is used. For example, a researcher may need to fit the PRF model multiple times as changes are made to a pre-processing pipeline which precedes the model; however, due to time demands, the researcher testing such changes to the pipeline may need to restrict the number of variations tested. Others have used a simplified PRF model to speed up the computation (Benson et al., 2018). Many researchers might be freed from such limitations by a method which accelerates the PRF estimation procedure.

Here, we present a software system called “qPRF” for fitting the PRF model that speeds up the underlying computations by a factor of 1436 when tested on a benchmark described herein. The system achieves this level of acceleration by making use of a searchable tree-like data structure. Each node of the data structure contains a parameter combination as well as the pre-computed predictions of the PRF model for that parameter combination. Nodes are related in the tree on the basis of similarity, allowing the system to search for an optimal parameter combination quickly. Next, we provide a detailed description of the qPRF and characterize its performance. Finally, we will discuss some potential applications and variants of this new method.

## 2. Methods

### 2.1. The PRF model

In human retinotopic mapping, the blood-oxygenation-level-dependent (BOLD) signal yielded by fMRI is the primary measurement of interest. As shown in Fig. 1, the BOLD signal (red) is a time series, which we shall denote *β*(*t*). This time series is associated with a single point within the cortex (usually a vertex of a mesh representing the cortical surface, but also sometimes a voxel within the cranial volume). The BOLD signal peaks when the neural activity in the given region is most active. The goal of PRF modeling is to identify a region within the visual field where the pattern of stimulation most accurately predicts the observed BOLD signal.

**Figure 1:**
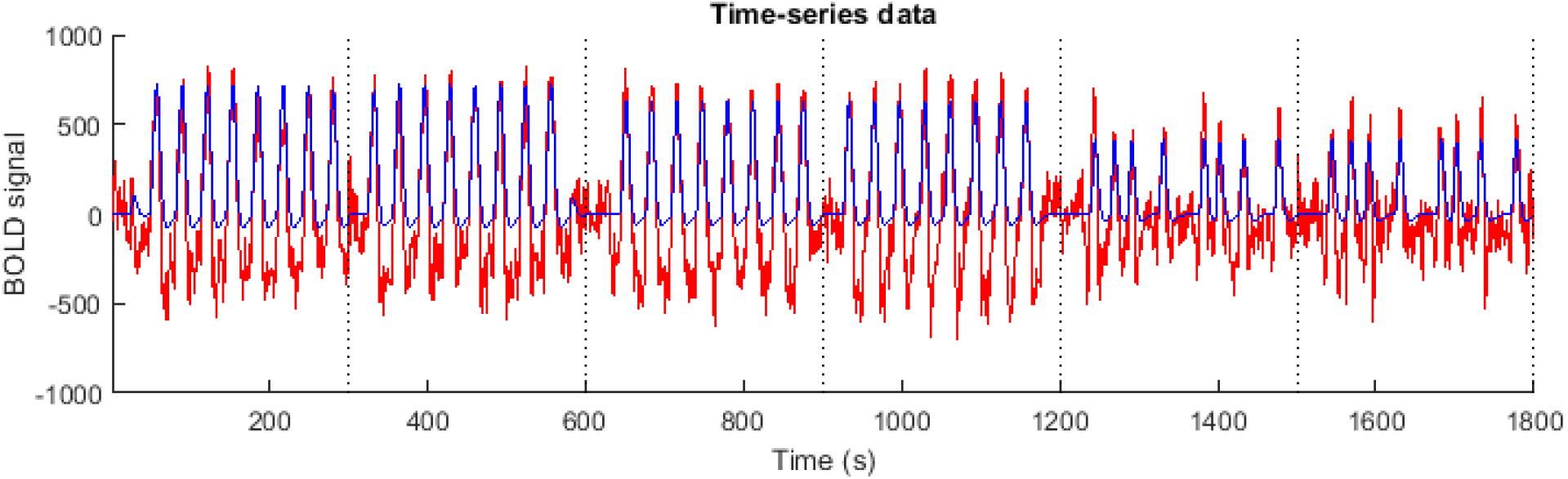
Visualization of BOLD signal (red) at a single point within the visual cortex of a subject and the prediction of the best-fitting PRF model (blue). Dotted lines demarcate the beginning and end of different stimulus runs (six total, each 300 seconds in duration).

Let us consider the following formulation of the PRF model.

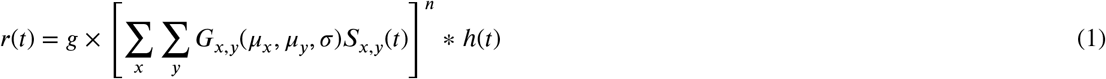

The output quantity, *r*(*t*), is shown in Fig. 1 (blue), and it is intended to match the BOLD signal as closely as possible. takes the stimulus (*S*_*x,y*_(*t*) which is an image indexed by *x* and *y* that varies in time *t*), uses a circular 2-D Gaussian receptive field (*G*_*x,y*_ parameterized by *µ*_*x*_, *µ*_*y*_, *and σ*) to extract a weighted sub-region of this stimulus, and integrates the activity within this sub-region using a compressive non-linearity (imposed by *n*), a hemodynamic response function (*h*(*t*) reflecting the general rise and decay times of neural activity upon stimulation), and a fixed gain factor (*g*). This yields the predicted population response, *r*(*t*). Eq. 1 explicitly denotes the five parameters of the PRF model: *g, µ*_*x*_, *µ*_*y*_, σ and *n*.

In PRF decoding, these parameters are estimated by choosing their values to minimize the sum of squared deviations between the model and the signal, ∑_*t*_(*β*(*t*)−*r*(*t*))_2_, and the current report presents a method designed to reduce the time demand of performing this estimation. The method addresses a processing bottleneck that is revealed in the following pseudo-code for a typical estimation procedure:

#### Algorithm 1

Generic PRF estimation procedure

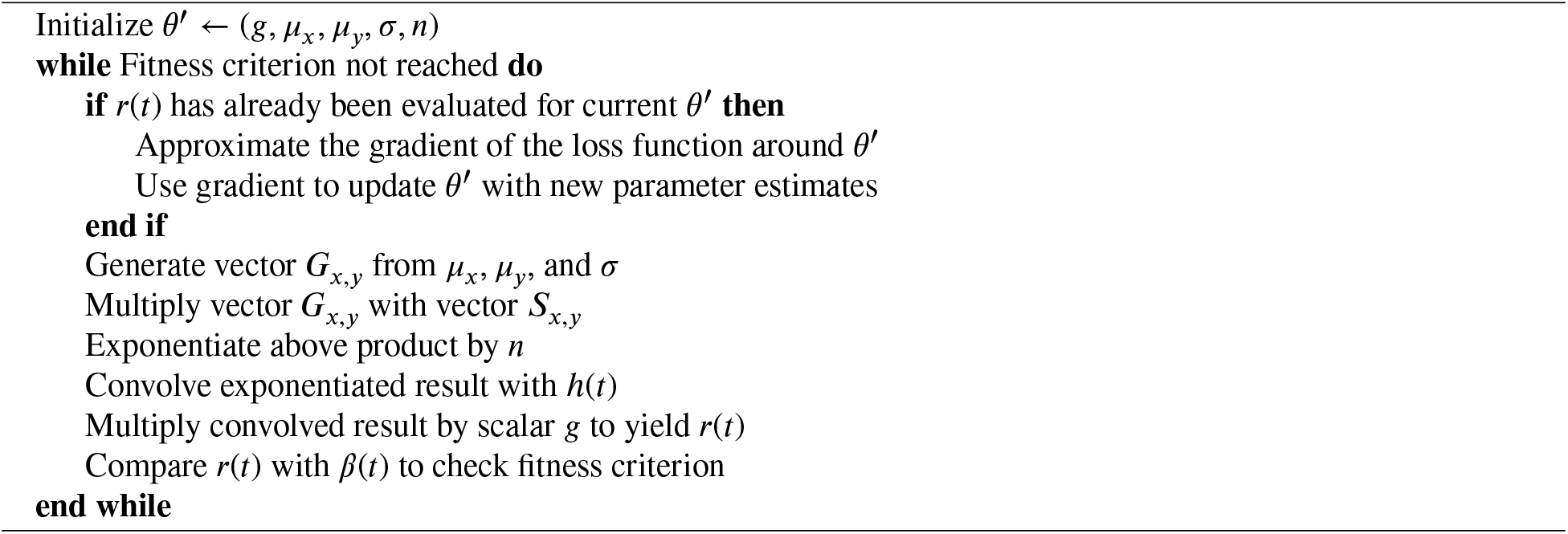

There are two critical observations regarding this pseudo-code. First, observe that several computations must be performed to simply evaluate *r*(*t*). Each of these computations introduces a time cost, especially those steps related to vector operations (i.e., multiplication of the 2D receptive field *G*_*x,y*_(*µ*_*x*_, *µ*_*y*_, σ) with the stimulus movie *S*_*x,y*_(*t*)) and convolution (i.e., convolution of the output of the receptive field by the hemodynamic response function *h*(*t*)). These steps must be performed on every iteration of the estimation procedure, thus slowing the total estimation time over the course of many iterations. A key insight behind the development in the current report is that these steps can be approximated by a single query of a data structure stored in memory.

Second, observe that the estimation procedure relies on a gradient method to search the parameter space. The approximation of the gradient is a similarly costly operation, requiring the PRF model to be evaluated at multiple parameter combinations around *θ*^′^ (in essence, requiring additional iterations of the code highlighted above for evaluating *r*(*t*)). Moreover, the PRF model is nonlinear, allowing local minima to exist in the objective function, so a gradient-based parameter search may be inefficient because the search algorithm may converge on a sub-optimal local minimum. In the case of analyzePRF, a standard package for PRF decoding (Kay et al., 2013), the gradient method used is the Levenberg-Marquardt algorithm, which is known to be susceptible to such convergence issues. Further, assuming the gradient is already known, this algorithm requires a matrix to be formed from the gradient vector computed around *θ*^′^ and requires this matrix to be inverted. Although these operations involve relatively small matrices (5 × 5, with sides corresponding to the number of parameters), a single vertex may require up to 500 iterations of this algorithm in analyzePRF, so the accumulation of these matrix operations may slow down PRF decoding non-negligibily. The second key insight of the work reported here is that search can be made more efficient by directing the fitting algorithm to pre-computed candidates that have been organized on the basis of similarity rather than having the fitting algorithm search for candidates naively.

### 2.2. Anticipatory modeling

Note that the steps contained in the while-loop of Algorithm 1 are conditioned only on knowing *S*_*x,y*_(*t*) and having a given parameter estimate *θ*. Moreover, for a given retinotopy experiment, *S*_*x,y*_(*t*) is typically known and fixed. Thus, in principle, we may pre-compute *r*(*t*) for all parameter combinations *θ* in a suitably large set Θ, anticipating the parameter combinations which might be yielded by PRF modeling. This “anticipatory” modeling may be done even before data collection has been initiated. Moreover, as is evident in Fig. 1, the BOLD signal which is to be modeled is often contaminated by substantial amounts of noise. Thus, our set of pre-evaluated parameter combinations Θ does not have to be very dense to achieve an adequate fit given typical levels of measurement noise. Further, we know that the parameter space to be considered has a number of boundaries and empirically established trends. For example, the gain parameter *g* is typically assumed to by non-negative, the values of *µ*_*x*_ and *µ*_*y*_ should be reasonably close to the stimulus location in the visual field (if not strictly within the stimulus location), and σ increases linearly with distance of the receptive field center from the point of fixation (Dumoulin and Wandell, 2008).

In the anticipatory modeling stage of our method, the stimuli are input to a program which evaluates the PRF model based on these stimuli for a large representative set of parameter combinations, Θ. The time series yielded by the PRF model evaluated at a given *θ* ∈ Θ, which we shall denote *r*(*t*|*θ*), is stored permanently in memory and associated with *θ*. By storing these results in memory, we no longer need to search within Θ directly for optimal parameters; rather, we can search the set of recorded *r*(*t*|*θ*) for a solution which minimizes the loss function ∑_*t*_(*β*(*t*) − *r*(*t*|*θ*))^2^.

As we shall describe below, the values of *r*(*t*|*θ*) are stored relationally, with similar values being linked together. This means that, once a roughly well-fitting candidate *r*(*t*|*θ*) is found, its relatives can be searched for a better fitting *r*(*t*|*θ*^′^). This kind of relational search can be repeated a number of times until an optimal fit is found. Since all candidate *r*(*t*|*θ*) values in this linked data structure have been pre-computed, the work of fitting is reduced to traversing the data structure and testing only the selected candidates against *β*(*t*).

### 2.3. Searchable tree structure

Given that we choose Θ to represent all of the reasonable parameter combinations that the given stimulus *S* might yield, Θ is, in general, too large to search exhaustively, so the recorded values of *r*(*t*|*θ*) for all *θ* ∈ Θ are stored in a rapidly searchable tree structure, represented schematically in Fig. 2. To reduce the number of comparisons, we organize this data structure on the basis of similarity, selecting several prototype *r*(*t*|*θ*) values to be searched initially at the coarsest level. These prototypes comprise a relatively small set of examples *r*(*t*|*θ*) values, reflecting, as completely and concisely as possible, all of the distinct variations of *r*(*t*|*θ*) achievable by the PRF model. Note that in Fig. 2, the top-most layer which represents these prototypes contains parameter combinations which are more sparsely distributed around the visual field than any layer below it, with the exception of a small central region (*ρ* < 0.4°) near the center of the visual field. Each parameter combination at the prototype layer points to a partition of the layer below it, representing all the parameter combinations within the layer that are near the prototype in their location on the visual field; members of the central region, however, point directly to the lowest layer because they already densely cover the visual field of their vicinity. The clustering of the secondary layer to members of the prototype layer on the basis of visual field location represents a very intuitive notion of similarity since visual field location has great influence on the shape of *r*(*t*|*θ*). Each parameter combination in the secondary layer subsequently points to a partition of the lowest layer, which represents variations of the parameters *n* and σ at the visual field location given by its parent. The colored paths in Fig. 2 trace how parameter combinations are selected at each level, leading to an optimized parameter combination at the lowest level.

**Figure 2:**
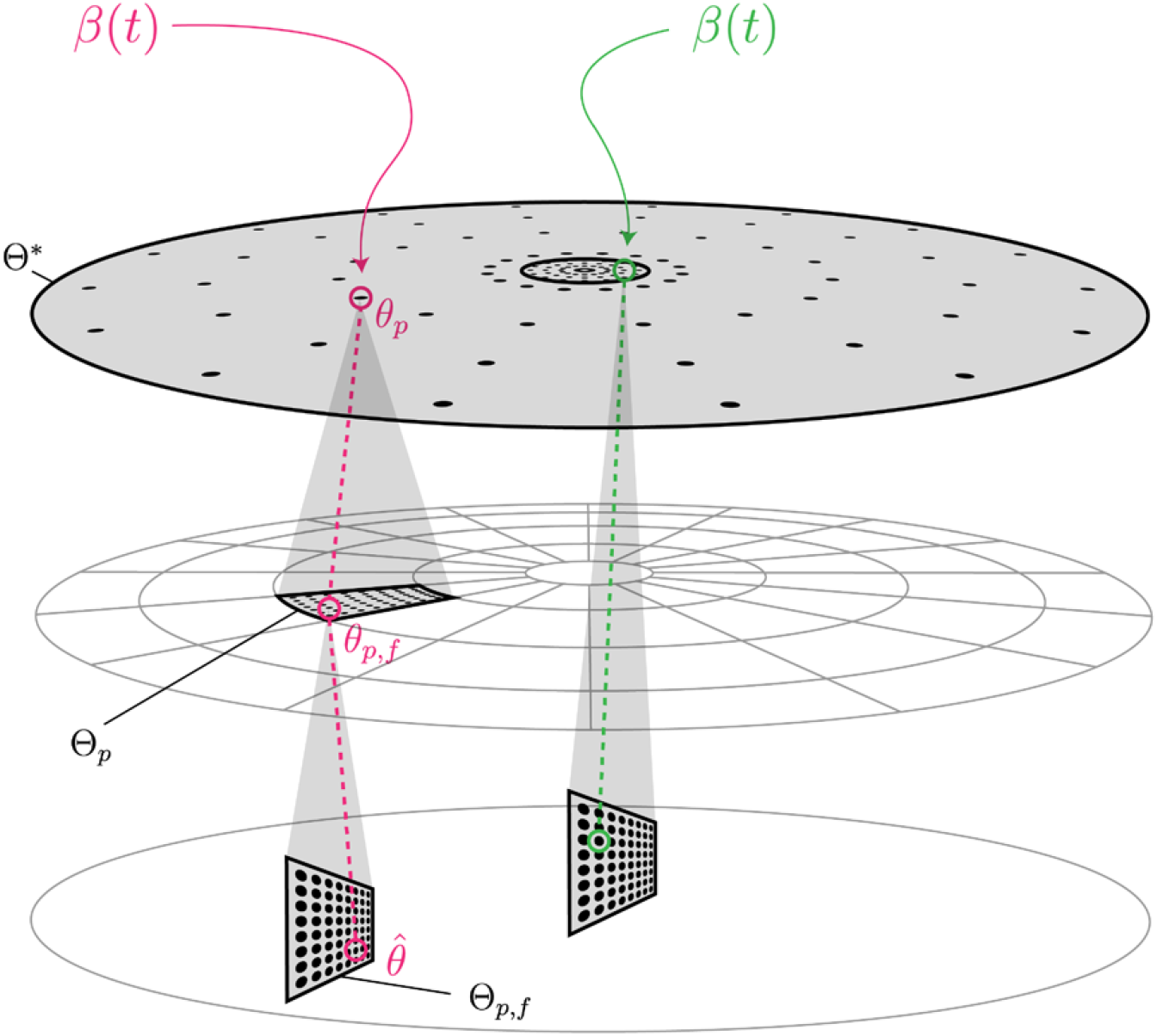
Diagram representing the order in which parameter combinations are searched within the tree-like data structure of qPRF when fitting a BOLD signal. Each black dot represents a single parameter combination. Each disk represents one of the three layers of the tree structure, as well as the visual field in which the parameter combinations are embedded. The green path shows an example search path for an observed BOLD signal *β*(*t*) which is best fit in the coarsest layer to a prototype within the central region, and the pink path shows an example search path for an observed BOLD signal *β*(*t*) which is best fit in the coarsest layer to a prototype *outside* the central region. Shaded regions encompass all of the parameter combinations which must be searched in the course of finding an optimal parameter combination. Note that parameter combinations are arranged orthogonally to the disk at the lowest layer because all the parameter combinations within a partition of this layer share the same visual field location and differ only in *n* and σ.

Let us say that *θ*_*p*_ is the parameter vector of prototype *p*, where the set of parameter vectors *θ*_*p*_ for all *p* form a set Θ^*^ ⊂ Θ. The first step of searching the data structure is thus to compare *β*(*t*) with *r*(*t*|*θ*_*p*_) for all prototypes *θ*_*p*_ ∈ Θ^*^ to find the best fitting member of this subset.

The search proceeds similarly at the finer levels of the data structure. Each prototype *p* points to a subset Θ_*p*_ ⊂ Θ, where Θ_*i*_ ∩ Θ_*j*_ = ∅ for *i* ≠ *j*. Once a best fitting prototype *p* is found at the coarsest level, Θ^*^, only its child subset Θ_*p*_ is searched for a better fitting member, *r*(*t*|*θ*_*p,f*_), where *θ*_*p,f*_ ∈ Θ_*p*_. For prototypes whose elements *µ*_*x*_ and *µ*_*y*_ are within a small radius of the point of fixation (the central region), Θ_*p*_ = {*θ*_*p*_}, allowing this secondary search to be skipped. This is because, in practice, the members of the set Θ^*^ differ primarily in *µ*_*x*_ and *µ*_*y*_, and are chosen to densely sample *µ*_*x*_ and *µ*_*y*_ within the central region. Prototypes with *θ*_*p*_ that fall outside the region are distributed more sparsely, and this secondary search is performed primarily to enhance the precision on *µ*_*x*_ and *µ*_*y*_ in this outer region.

The third and final level of the data structure only represents variation in σ and *n*. Once a best-fitting *θ*_*p,f*_ ∈ Θ_*p*_ is found, its child subset Θ_*p,f*_ ⊂ Θ is searched to optimize σ and *n*. The optimal parameter combination found within Θ_*p,f*_ represents the final estimate which is returned by qPRF, which we denote 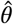. Note that, given *µ*_*x*_, *µ*_*y*_, σ, and *n*, the value of *g* which minimizes the sum of squared deviations can be calculated directly, so *g* in our program is not a dimension of Θ. Further, note that at the two coarsest levels of the tree, the σ value used among candidates is a linear function of the distance between *µ*_*x*_ and *µ*_*y*_ from the fixation point (reflecting the known positive relationship between visual field eccentricity and receptive field size; Smith, Singh, Williams and Greenlee, 2001; Dumoulin and Wandell, 2008). Altogether, the data structure resembles a tree, shown in Fig. 2, where the first two levels facilitate a two-phase search for an optimal receptive field location, and the third level facilitates optimization of the receptive field size and compressive non-linearity given this location.

For the test implementation examined below, we use a total of 1,104,960 parameter combinations. The coarsest level of the data structure, Θ^*^, comprises 552 of these combinations, which we call prototypes. These prototypes are distinguished primarily by their values of *µ*_*x*_ and *µ*_*y*_. Since we choose the values of *µ*_*x*_ and *µ*_*y*_ to evenly cover the visual field in polar coordinates, let us instead use the polar parameterization *ρ* and ϕ to refer to these parameters hereafter. The central region, a circle of radius 0.4° at the center of the visual field, contains 264 of the parameter combinations in Θ^*^, representing all combinations of 6 values of *ρ* (evenly spaced from 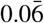 to 0.4) and 44 values of ϕ (evenly spaced from 0° to 360°, starting at 0° and ending at 351.8°). For all the central protoypes, *n* = 0.1, and σ was an increasing function of *ρ* (described below). As noted above, if the best fitting prototype in Θ^*^ resides within this central region, the search short circuits and skips over searching Θ_*p*_.

In contrast to the central prototypes, the prototypes outside the central region are distributed more sparsely in *ρ* and ϕ because, outside the central region of the Θ^*^, all of the prototypes *do* point to a corresponding Θ_*p*_ with densely packed *ρ* and ϕ values which is subsequently searched. The exterior of the central region in Θ^*^ is itself divided into two regions, which we refer to as the *para-central* and *peripheral* regions and which are separated by a circle of radius 8° in the visual field (the outer edge of the stimuli used in the HCP dataset). In the para-central region, there were 160 prototypes, corresponding to all combinations of 10 values of *ρ* (evenly spaced within the para-central region spanning 0.4° to 8°) and 16 values of ϕ (evenly spaced, starting at 0°). In the peripheral region, there were 128 prototypes representing 8 values of *ρ* (evenly spaced in the peripheral region spanning 8° to 16°) and 16 values of ϕ. Across all prototypes, the function used to specify σ was

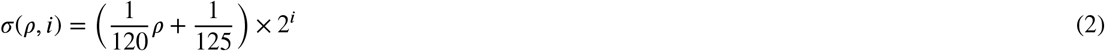

where the units of σ and *ρ* are both given in degrees of visual angle. At the lowest level of the data structure, the values of *i* used to specify the values of σ to be searched at any given value of *ρ* were the integers 1 through 8. For central and para-central prototypes, the value of *i* was chosen to be 4, and for peripheral prototypes, the value of *i* was chosen to be 5. The values of *i* used among prototypes were chosen to reflect the following known properties of receptive fields: (1) receptive field size near the fovea is markedly smaller than elsewhere in the visual field, and (2) outside the foveal region, the slope of the linear trend in receptive field size appears to increase multiplicatively with higher-order visual areas (i.e., V2 and V3, as shown by Dumoulin and Wandell, 2008). The range *i* ∈ {1, ⋯, 8} captures the wide variety of σ values observed across these areas.

All para-central and peripheral prototypes pointed to a child parameter space Θ_*p*_ containing 95 combinations to be subsequently searched. Thus, the total number of parameter combinations contained in the second level of Fig. 2 equals 27,360 (288 non-central parent prototypes × 95 combinations in each child parameter space). The 95 children of each non-central prototype represent combinations of 5 values of *ρ* and 19 values of ϕ. These values of *ρ* and ϕ were centered on the *ρ* and ϕ values of the parent prototype and were chosen so that the values of *ρ* and ϕ formed a uniform grid when considering all members of the second level in Fig. 2 (with a change in the grid resolution when going from the para-central to peripheral region, corresponding to a fixed number of children but a reduced number of parent prototypes).

All 27,360 children of the second level of the data structure, as well as all 264 central prototypes, each pointed to a unique parameter space Θ_*p,f*_ which shared the values of *ρ* and ϕ as its parent and comprised 40 variations of *n* (5 levels) and σ (8 levels). Across all Θ_*p,f*_, the values of *n* were 0.025, 0.05, 0.1, 0.2, and 0.4. The 8 values of σ represented within a given Θ_*p,f*_ were derived from Eq. 2 using the *ρ* value of the parent, and 8 evenly spaced values of *i* going from −4 to 4. Thus, at the finest level of the data structure, we have (27, 360 + 264) × 40 = 1, 104, 960 parameter combinations. Although this parameter space would be too massive to search exhaustively under reasonable time constraints, the parent-child relationships represented by the structure make it so that, altogether, a vertex which is mapped to a central prototype will require only 552 +40 = 592 comparisons in the course of estimation, and a vertex mapped to a para-central or peripheral prototype will require 552 + 95 + 40 = 687 comparisons.

### 2.4. qPRF algorithm

The following pseudocode describes the fitting procedure performed by qPRF.

#### Algorithm 2

qPRF estimation procedure

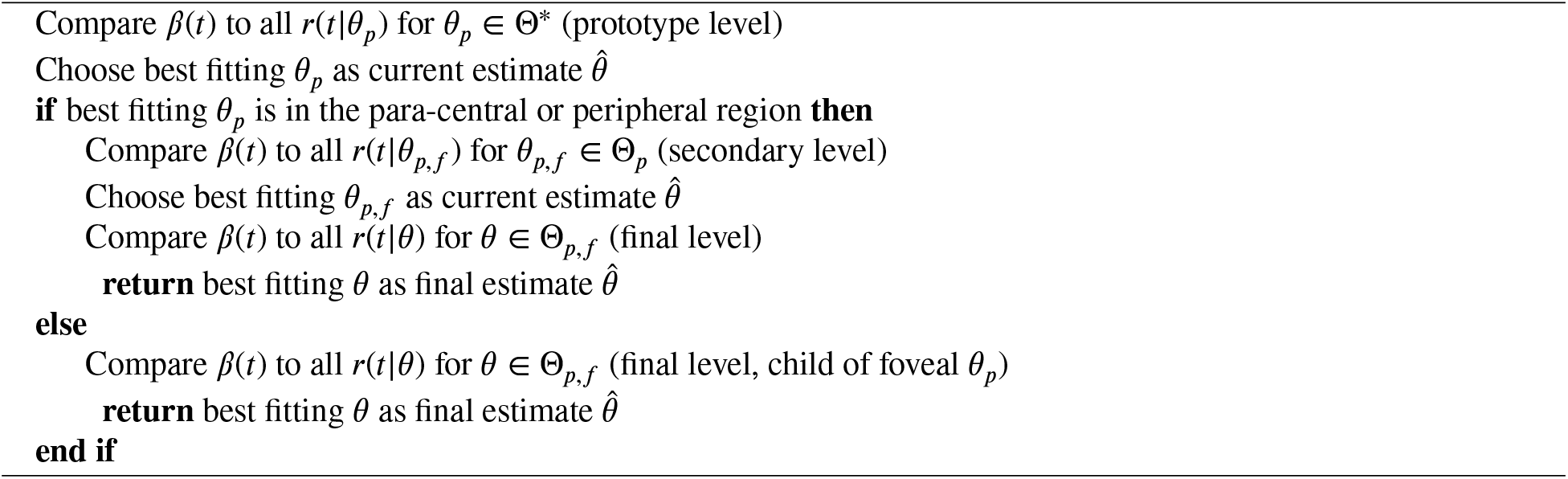

## 3. Results

The data used in this analysis were originally sourced from the Human Connectome Project (HCP; Ugurbil et al, 2013; Van Essen et al., 2013) 7T Retinotopy Dataset. This dataset represents the largest publicly available collection of brain imaging data from human subjects in a retinotopy experiment. Complete details of this experiment were reported by Benson et al. (2018), who also provided PRF estimates based on these data. The PRF estimates computed by Benson et al. are available online (https://osf.io/bw9ec) and served as a reference set of estimates which we compare our qPRF estimates to.

The HCP dataset includes 181 subjects aged 22 to 35. For each subject, a total of 91,282 vertices were decoded with the PRF, covering both hemispheres of the cortex and subcortical regions. Of these, 29,696 (29,716) vertices represented the surface mesh of each subject’s left (right) hemisphere. Thus, altogether, the dataset represents 10,753,572 unique vertices, each of which presents a single opportunity to perform PRF estimation. The cumulative BOLD signal at each vertex was a vector of 1,800 time points, comprising six runs of 300 time points (frames), with a repetition time (TR) of 1 second per frame.

For the HCP study, the time required to generate the data structure underlying the qPRF computation was 4.01 hours on a 3.50GHz Intel Xeon E5-1650 v3 CPU. The total memory required to store the data structure was 6.1 GB.

The total time required to the generate qPRF estimates for the entire HCP data set was 15.2 hours on a 3.50GHz Intel Xeon E5-1650 v3 CPU. When compared to another widely available PRF decoder, analyzePRF (Kay et al., 2013), estimation via qPRF reduced the computation time by a factor of 1436. The comparison used to derive this factor is shown in Fig. 3. Note that, for this comparison, we did not re-run estimation on the entire data set via analyzePRF as this would be too time intensive. Instead, we ran analyzePRF and qPRF on a subset of 100 vertices from each hemisphere of 32 subjects. Whereas analyzePRF required 13.6 hours to complete this analysis, qPRF required only 34.5 seconds. According to this benchmark, qPRF takes 5.4 milliseconds on average to decode a single BOLD signal, whereas analyzePRF takes 7.7 seconds.

**Figure 3:**
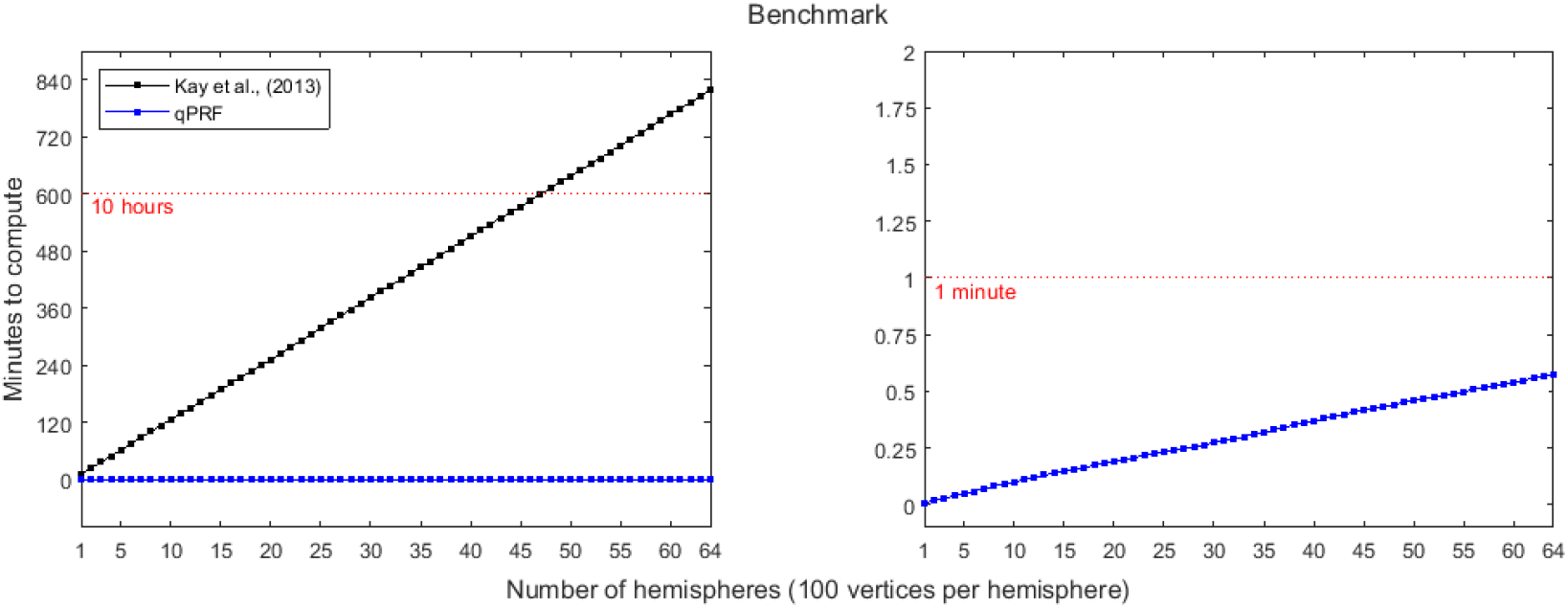
Comparison between qPRF and analyzePRF (Kay et al., 2013) in terms of time to compute estimates for the PRF model. Note that left and right panels visualize the same information. The *y*-axis of the right panel has been scaled to make the increasing computation time of qPRF visible. Since analyzePRF required 12.5 minutes to compute a single hemisphere, the analyzePRF time curve is not visible in the right panel.

There are three important distinctions to make as part of this comparison. (1) Whereas qPRF uses a novel optimization procedure specified above, analyzePRF makes use of the Levenberg–Marquardt algorithm, available in Matlab as the function **lsqcurvefit.m**. For each vertex, analyzePRF performed up to 500 iterations of the Levenberg-Marquardt algorithm, returning sooner if the change in the objective function or parameter estimate was smaller than 1×10−6. (2) Our benchmark made use of parallel computation to speed up analyzePRF, a feature which is provided with the analyzePRF package; by contrast, the qPRF computations were performed serially due to the memory limitations of our benchmarking machine. In principle, even greater reductions in computation time may be achieved by distributing the workload to multiple qPRF workers. (3) Below, we use results from Benson et al. (2018) to compare the quality of qPRF estimates to a standard set of PRF estimates. Benson et al. made use of a different version of analyzePRF than the one used in our benchmark. Namely, they used a fixed value of *n* = 0.05 whereas we estimated all 5 parameters of the PRF model in Eq. 1. This likely sped up the computation time for Benson et al. since the version of analyzePRF tested here performs a two-stage estimation procedure wherein all parameters except *n* are first optimized under a fixed *n*, then all parameters including *n* are optimized simultaneously. Benson et al. needed only to perform the first stage of this optimization procedure to form their estimates.

The qPRF estimates yielded *R*^2^ measures that were markedly similar to those reported by Benson et al. (2018). Like Benson et al. (and Kay et al., 2013, similarly), we define *R*^2^ to be a percentage calculated as

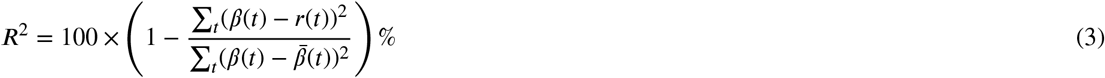

where 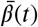 is the mean of the observed BOLD signal *β*(*t*) over time. Fig. 4 shows a heat map of the difference between *R*^2^ values achieved by Benson et al. 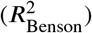 and those achieved by the qPRF model 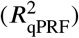 across all vertices of both hemispheres of all subjects in the HCP dataset. We computed 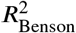 by regenerating the BOLD signal predicted by Benson et al.‘s PRF estimates and computing *R*^2^ between the predicted and observed signals. Units of *R*^2^ are reported as percentages ranging from 0% to 100%. The color of the heat map represents the number of vertices within the HCP dataset for which the 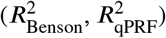-coordinate fell within the corresponding 1% × 1% bin. The main observation to make regarding Fig. 4 is that the peak of the heat map is a ridge that closely follows the positive diagonal of the heat map, indicating equivalence between the two solutions in terms of goodness-of-fit. The distribution of vertices around this line is skewed positively, indicating that some of the qPRF estimates were better fitting than their corresponding estimates from Benson et al.. The middle 95% of differences 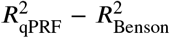 fell within the interval [0.02, 5.51], with a median of 0.39 (reported here as the absolute difference between the two percentages). Although 99.7% of vertices were more closely fit using qPRF, the above interval indicates that these differences were, in many cases, not practically significant. Although we did not re-produce their PRF estimates ourselves, Benson et al. noted that their provided estimates were based on a modified version of analyzePRF. Most notably, Benson et al. restricted *n* = 0.05. Instead, we estimated the exponent parameter, *n*. Discrepancies between our solutions and theirs might be attributed to this difference. Nonetheless, note that when the PRF estimates from Benson and from qPRF are plotted side-by-side on the cortical mesh (Fig. 5), the results look strikingly similar. As cited earlier, such plots are typically used as diagnostic images upon which experts may subjectively identify the boundaries of the visual areas (e.g., Dougherty et al., 2003, Benson et al., 2022, Himmelberg et al., 2023), so Fig. 5 may be taken to suggest that qPRF and analyzePRF offer comparable results to the end user, except that qPRF yields results in a shorter time.

**Figure 4:**
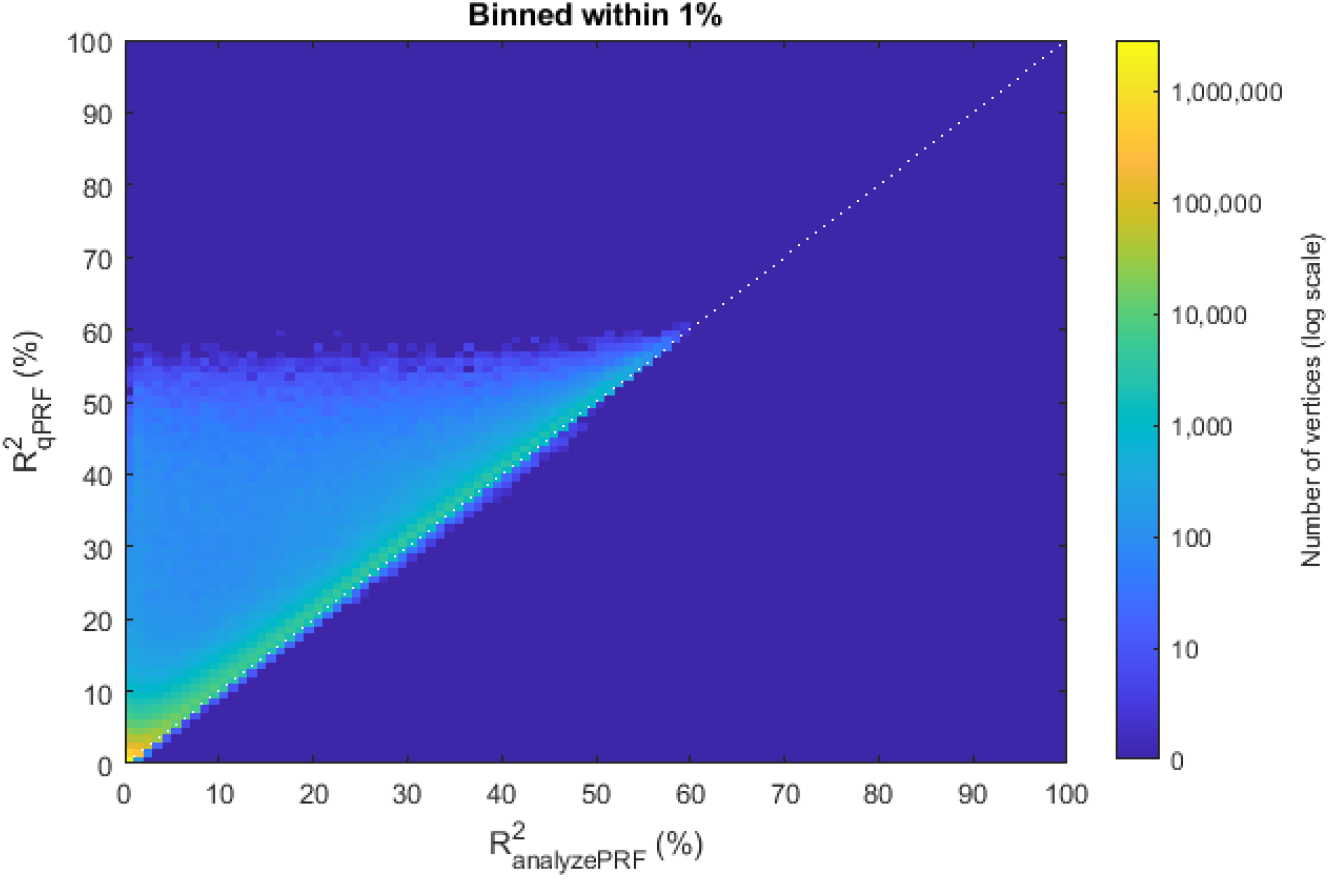
Heat map comparing the *R*^2^ values of the solutions achieved by qPRF against the solutions provided by Benson et al. (2018). Counts are binned within 1% and are shown on a logarithmic color scale. The dotted white line indicates equivalence of the two solutions in terms of *R*^2^. The peak of this distribution follows this line, indicating that the goodness-of-fit of the two solutions was strikingly similar; however, the distribution has a long tail skewing in the direction of the qPRF solutions, which achieved a greater *R*^2^ than the solutions provided by Benson et al. on 99.7% of vertices. The median difference was very small (0.39%, where *R*^2^ can range from 0% to 100%). Note that the vast majority of vertices yield an *R*^2^ near zero for both solutions, reflecting a lack of PRF information in the BOLD signal at most vertices of the brain mesh.

**Figure 5:**
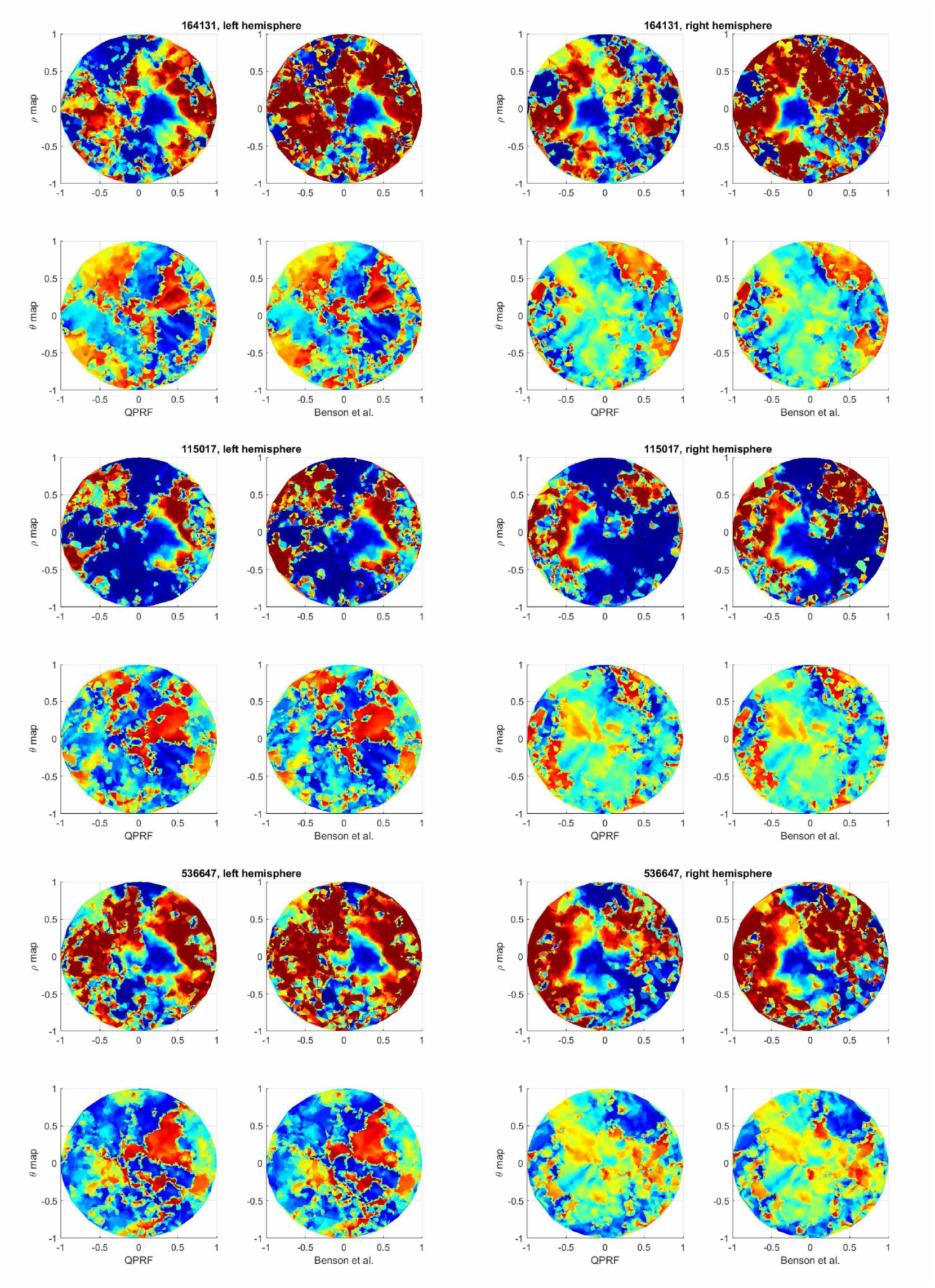
Comparison of the PRF solutions yielded by qPRF and those provided by Benson et al. (2018). The figure contains six panels; each row of panels represents one subject, and each column of panels represents one hemisphere. Within each panel, we see the *ρ* estimates and *θ* estimates yielded by qPRF and provided by Benson et al. projected conformally onto a parametric disk (Ta et al., 2022). The three subjects selected as examples are those also used as examples by Benson et al.. Note that all *θ* estimates share the same color scale as do all *ρ* estimates (which are clipped to a single color for *ρ* > 8°, the maximum extent of the stimuli). The *θ* estimates are strikingly similar. The *ρ* estimates share much of the same global structure, with the solutions from Benson et al. having more extreme estimates in some regions.

## 4. Discussion

The PRF model formulated by Dumoulin and Wandell (2008) has been an essential tool in visual neuroscience. The model addresses the basic challenge of retinotopic mapping, which is to identify a region in the visual field that corresponds to each vertex on the visual cortex. Its application is of foundational importance to studying visuocortical function, providing experts with decoded images of fMRI upon which they can manually identify the boundaries of visual areas (e.g., Benson et al., 2022; Himmelberg et al., 2023). Such decoding is not only essential to studying the functional organization of the brain, but may prove to have a wide variety of clinical applications, e.g., in the tracking of AD (Brewer and Barton, 2014; Brewer et al., 2016), glaucoma (Duncan et al., 2007; Boucard et al., 2009), MD (Bridge, 2011; Baker et al., 2005), stroke (Elshout et al., 2021), or in measuring the effect of noninvasive treatments for such conditions, like VRT (Barbot et al., 2021).

In this study, we attempted to address a technical challenge in fitting the PRF model to fMRI retinotopy data. We have reported the details of a novel method of PRF model estimation, which we call qPRF, to reduce the time required to decode fMRI. Although the technique requires some initial pre-processing (approximately 4 hours for the parameter space defined here) and also requires some static memory consumption (approximately 6 GB in our example usage), the time efficiency gained outweighs the relatively small upfront resource demands. In our benchmark, we found that qPRF was able to provide estimates within 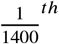 the time required for another popular package (analyzePRF) to accomplish the same task.

The qPRF achieves its accelerated operation time by making use of a searchable tree structure which stores parameter combinations and corresponding prediction curves of the PRF model given a stimulus set that is fixed across subjects. Although in our analysis of the qPRF method’s performance, we considered only the particular stimuli used in the HCP retinotopy dataset (Benson et al., 2018), our description of the qPRF method does not assume a specific stimulus *S*_*x,y*_(*t*) (Eq. 1) and is general to other kinds of visual stimuli for which the PRF model itself applies. Indeed, the principles which underlie the qPRF method’s time cost reduction (i.e., anticipatory modeling and relational storage of pre-computed predictions) are not specific only to retinotopic mapping and may be applied, for example, to tonotopic or somatotopic mapping in an intuitive way; of course, the exact details of Eq. 1 will need to be altered to reflect the forms of the corresponding perceptual signals and receptive fields in such extensions of the method. The qPRF method may also offer advantages when incorporated into existing techniques for constraining and characterizing an ensemble of PRF estimates (e.g, Tu, Ta, Lu and Wang, 2021; Xiong, Tu, Lu and Wang, 2023).

In the current implementation of the qPRF, the design and sampling density of the data structure were based on our understanding of the range of parameters and measurement noise in retinotopic mapping experiments. However, we acknowledge that these parameter ranges may still not be optimal for every application and may be further optimized. For example, some users may want to increase the sampling density to achieve more precise solutions. Following Benson et al. (2018), fixing *n* might further speed up the algorithm by reducing the number of parameter combinations that must be modeled in anticipation and subsequently searched. It is also worth noting that although the hemodynamic response function, denoted *h*(*t*) in Eq. 1, has been treated here as a constant, it has a parametric form, and in principle might be optimized within the framework of qPRF by introducing additional free parameters. In this variant of qPRF, the principles of anticipatory modeling and relational storage would still apply, but the variety of prediction curves *r*(*t*) would be more diverse, reflecting the alternate configurations of *h*(*t*). Some of these configurations might be able to capture large negative dips in the BOLD signal, as seen in Fig. 1, which the specific implementation of qPRF described here (like standard implementations of the PRF model) does not capture.

Now that we have shown that the standard PRF model can be estimated rapidly with qPRF, we can build on this framework to estimate more elaborate forms of the PRF model under similarly reduced time costs. We have already shown that qPRF allowed us to estimate a value for the exponent parameter, *n*, of the PRF model whereas in past literature, this was assumed to be fixed for simplicity (Benson et al., 2018). An elaboration of the PRF model where the techniques of qPRF may be applied is one where the receptive field is non-circular. In this case, the receptive field of a particular voxel on the visual field may be elliptically elongated with some orientation. Although implementations of such a model may vary, this generalization introduces at least two additional parameters which must be estimated. In principle, this elliptical qPRF can be based on the existing qPRF data structure described here, with one additional layer of parameter combinations added at the finest level to capture the additional variations in the receptive field shape. Although standard packages for PRF decoding do not consider this possibility, there are well-known perceptual phenomena, such as visual crowding (Whitney and Levi, 2011), wherein the perceptibility of a visual feature depends on how the feature is oriented in the visual field periphery, suggesting that the receptive field is elongated orthogonal to this orientation. Other elaborations on receptive field structure include center-surround weighting functions, where stimulation can have both excitatory and inhibitory influences on the predicted signal depending on its distance from the receptive field center. Such an elaboration on the PRF model might also be able to capture the negative dips in the observed BOLD fMRI signals which are otherwise unexplained by the standard PRF model.

The search algorithm itself can also be changed based on user defined needs. In the implementation of qPRF detailed here, we describe a unidirectional search path, considering only the single optimal candidate at each level of the data structure. However, the data structure provided by qPRF can facilitate other search methods. For example, the search may consider the top 5 or 10 candidates at each level and all of their children. Although this increases the number of comparisons to be performed, the number of candidates to be considered in parallel can be tuned to the researcher’s competing needs for precision and time efficiency. This alternative approach might be helpful in cases where, conditioned on a new estimate of *n* and/or σ, the optimal estimates of *µ*_*x*_ and *µ*_*y*_ change. Other search methods that the qPRF data structure can support include stochastic optimization, where the children of a small number of random non-optimal candidates are searched as a security against optimizing to a local minimum. In our view, a variety of techniques might be developed using the basic principles of the qPRF method to enhance PRF modeling, a tool which has already proven to be uniquely important in visual neuroscience.

In conclusion, we have introduced qPRF, a novel software system for population receptive field (PRF) decoding of BOLD fMRI. This system operates at speeds about 1400 times faster than traditional PRF decoding systems, while delivering parameter estimates that match those obtained with conventional methods. The qPRF may facilitate the development and application of advanced models for decoding sensory maps, potentially advancing our understanding of the human brain.

## Code availability

The qPRF software, with example code to generate the searchable tree structure and to fit the PRF model, is available here.

## Declaration of interests

Financial support was provided by National Eye Institute Grant R01 EY 032125. Authors S.W., Y.W., and Z.-L.L. have a provisional patent to New York University and Arizona State University, docket number 300727US01-109197-0000142.

## CRediT authorship contribution statement

**Sebastian Waz:** Algorithm development and evaluation; Manuscript writing. **Yalin Wang:** Manuscript revision. **Zhong-Lin Lu:** Project conceptualization; Algorithm design; Manuscript Revision.

## Notes

### Competing Interest Statement

The authors have intellectual property interests in the technology described in this paper.

